# MARS: Motif Assessment and Ranking Suite for transcription factor binding motifs

**DOI:** 10.1101/065615

**Authors:** Caleb Kipkurui Kibet, Philip Machanick

## Abstract

We describe MARS (Motif Assessment and Ranking Suite), a web-based suite of tools used to evaluate and rank PWM-based motifs. The increased number of learned motif models that are spread across databases and in different PWM formats, leading to a choice dilemma among the users, is our motivation. This increase has been driven by the difficulty of modelling transcription factor binding sites and the advance in high-throughput sequencing technologies at a continually reducing cost. Therefore, several experimental techniques have been developed resulting in diverse motif-finding algorithms and databases. We collate a wide variety of available motifs into a benchmark database, including the corresponding experimental ChIP-seq and PBM data obtained from ENCODE and UniPROBE databases, respectively. The implemented tools include: a data-independent consistency-based motif assessment and ranking (CB-MAR), which is based on the idea that ‘correct motifs’ are more similar to each other while incorrect motifs will differ from each other; and a scoring and classification-based algorithms, which rank binding models by their ability to discriminate sequences known to contain binding sites from those without. The CB-MAR and scoring techniques have a 0.86 and 0.73 median rank correlation using ChIP-seq and PBM respectively. Best motifs selected by CB-MAR achieve a mean AUC of 0.75, comparable to those ranked by held out data at 0.76 – this is based on ChIP-seq motif discovery using five algorithms on 110 transcription factors. We have demonstrated the benefit of this web server in motif choice and ranking, as well as in motif discovery. It can be accessed at http://www.bioinf.ict.ru.ac.za/.

## Introduction

We introduce MARS (Motif Assessment and Ranking Suite), a web server hosting a suite of tools for motif evaluation and ranking. It provides a service that is necessitated by the advance in high-throughput sequencing technologies at a continually reducing cost that has seen a large amount of – often noisy – data being generated by various studies [27]. To make sense of these datasets, several tools and algorithms have been developed, differing in data cleaning and statistical algorithms involved. The wide variety and the large number of computational tools being developed makes it hard for a non-specialist with limited computational skills to choose the best tools for use in their research [1]. Additionally, to improve on currently available tools, algorithm developers need well thought out and representative benchmark data (gold standard) and evaluation statistics. This problem has been tackled by independent evaluation studies [24,25,35,37] focused on various niches of research and data, producing incomparable results. This prompted a question of “who watches the watchmen” (evaluation benchmarks) by Iantorno *et al.*, [14] who also proposed that a proper benchmark should follow a set of pre-determined criteria to ensure that the evaluations are biologically relevant. Aniba *et al.*. [1] provide a set of criteria for a good benchmark that we adapt to the context of motif assessment as follows:

- **Relevant** – scoring methods should provide biologically meaningful evaluations
- **Solvable** – scoring methods should not be trivial but must be possible to use with reasonable effort
- **Scalable** – the benchmark should be expandable to cover new techniques and algorithms as they develop
- **Accessible** – data and statistical tools should be easy to source and use to evaluate other algorithms or protocols
- **Independent** – methods should not be tailored or biased to a particular algorithm or biased towards certain experimental techniques
- **Evolvable** – the benchmark should change as new data are made available, as well as to reflect the current problems and challenges in the field

The evaluation problem has been widely investigated in multiple sequence alignment, 3D structure prediction, protein function and gene expression analysis [1], mostly following the criteria above. However, evaluation remains an active challenge in gene regulatory research, especially in predicting TF binding sites and the accuracy of prediction models [39]. This difficulty is directly linked to the motif discovery problem, which has been attributed to the degeneracy of TF binding and the presence of multiple potential binding sites in the genome [13,16]. The difficulty of motif discovery has in turn driven the growth in experimental techniques developed to improve the affinity and specificity of TF binding site prediction models; techniques to identify binding sites or binding affinity include Chromatin Immunoprecipitation followed by parallel sequencing (ChIP-seq) [15] or exonuclease cleavage (ChIP-exo) [28], protein binding microarray (PBM) [2], Assay for Transposase-Accessible Chromatin with high-throughput sequencing (ATAC-seq) [7], DNase I digestion and high-throughput sequencing (DNase-seq) [31] and many others. Consequently, the number of algorithms and hence the binding models in databases continues to increase. Two areas are in need of evaluation: the algorithms used in motif discovery and the models deposited in the various motif databases. Although interlinked, in that ranking a model can be an indirect evaluation of an algorithm used to generate it, most of the evaluation attempts so far have been focused on the algorithms. This is a challenging task given that new tools are published regularly with varied implementations, scoring functions and even the data used for motif discovery. Therefore, establishing a widely useful model or motif algorithm evaluation platform is a moving target.

Nonetheless, there have been some attempts to develop tools and techniques to evaluate motif discovery algorithms, which can be categorized into assess-by-binding site prediction, motif comparison or by sequence scoring and classification [18]. We relate a selection of known approaches to the benchmark criteria we outline above.

An assess-by-binding site prediction approach evaluates algorithms by their ability to identify known or inserted binding sites in a sequence. It is widely investigated; a number of stand-alone motif assessment tools [26] and web servers [30,33] have been developed. However, these tools neither *evolved* nor *scaled* with advances in motif discovery algorithms, reducing their *relevance*, and thus failing to meet major requirements for an evaluation benchmark. Assess-by-scoring and classification tests binding models by their ability to discriminate sequences known to contain binding sites from those without. UniPROBE is the most comprehensive collection of PBM-derived motifs [22] and we are aware of one web server that uses such data in motif evaluation [36] (it has neither *evolved, scaled* not is it easily *accessible*) while, for ChIP-seq data, Swiss Bioinformatics hosts a simple web server (PWMTools (http://ccg.vital-it.ch/pwmtools/pwmeval.php) that is limited to testing single motifs against ENCODE data using sum occupancy scoring; it does not allow for comparative motif or sequence data testing (not *relevant* or *accessible*) and the site is not published. On the other hand, assess-by-motif-comparison has generally been used to determine if the discovered motifs are similar to those in ‘reference databases’ using motif comparison algorithms. An algorithm is considered successful if it can predict a motif similar to those in the database. However, this assumes the accuracy of previous predictions (not *relevant* or *scalable*), a weakness we address in this study.

The spread of motif models across DBs and in different PWM formats makes it difficult to create a benchmark that ranks multiple motifs for a given TF, and this problem is compounded by the growth in available data [18]. There is a lack of an easily *accessible* and *independent* motif evaluation platform that can allow users to rank PWM models for a given TF. To fill this gap, we introduce a web server that hosts a suite of motif assessment tools used to evaluate and rank motifs. For wider applicability, we collect ChIP-seq and PBM data generated from different labs and use an average score to represent a given motif, with the assumption that this would capture the most general binding behaviour. We also apply a wide variety of scoring functions and statistics to reduce technique bias. In addition, we introduce a novel Consistency-Based Motif Assessment and Ranking (CB-MAR) approach that can be considered to be *data-independent*, hence less biased compared to scoring-based techniques.

## Materials and Methods

### Benchmark data

We downloaded all ChIP-seq peaks uniformly processed by Analysis Working Group (AWG) from ENCODE [32], PBM from UniPROBE [5] database and PMW motifs from various databases and publications, prepared as previously described [18] and stored in a MySQL database (Table 1). Alternative TF names (from GeneCards [29]) link various alternative TF names to a TF class ID derived from the TFClass classification [38] to find all motifs for a TF irrespective of naming inconsistency. Unless otherwise specified, all sequence data used are derived form the human genome (hg19), except for the PBM data where we also use mouse data.

**Table 1.** Summary of Benchmark Data in the Database: ‘Used in analysis’ represent data using in our comparative tests. For motif discovery with GimmeMotifs we use 110 (in brackets).

| Data Type | TFs | motifs | motifs per TF | Used in Analysis |
| --- | --- | --- | --- | --- |
| ChIP-seq | 161 | 691 | 4.3 | 83 (110) |
| PBM | 285 | 455 | 1.6 | 60 |
| PWM-Motifs | 1352 | 6050 | 4.5 | 3686 |

#### Algorithms Overview

We have previously described the implementation of assessing by scoring and enrichment algorithms [18]. In summary, for each TF, the motifs in PWM format are used to score sequences partitioned into positive (test) and the negative (background) using one of the implemented scoring functions. Finally, the ability to classify the two sets is evaluated by area under receiver operator characteristic curve (AUC) or the mean normalized conditional probability (MNCP) statistics (Table 2). See our previous paper [18] for more details on the scoring functions and statistics used.

**Table 2.** Implemented scoring functions: The table provides a list of scoring functions used and the recommended statistics.

| Scoring functions | Preferred stat. | Recommended? | Reference |
| --- | --- | --- | --- |
| Energy | AUC or MNCP | High (Default) | [40] |
| GOMMER | MNCP | High | [8] |
| Sum Occupancy | MNCP | Average | [23] |
| Max Occupancy | MNCP | Average | [41] |
| Max Log-odds |  | Not Recommended | [41] |
| Sum Log-odds |  | Not Recommended | [3] |

### Consistency-Based Motif Assessment and Ranking (CB-MAR)

CB-MAR is based on the idea that ‘correct motifs’ are more similar to each other while incorrect motifs will differ from each other. The logic for this view is that differing methods are unlikely to reproduce each others’ errors. This idea is used in evaluating sequence alignments: correct ones are assumed to compare with each other in a consistent manner, while incorrect ones will differ from each other in various ways, generating inconsistent alignments [14,20].

We implement this approach using Tomtom [12] and FISim [11] motif comparison algorithms. For a given TF, we calculate a similarity score between all motif pairs and finally an average motif similarity score, which we use as a measure of motif quality. For best results, the benchmark motif set should be: (*a*) generated from a variety of data and motif finding algorithms – with (*b*) identical motifs eliminated (especially in a small set) – and (*c*) be large enough to capture variation in binding behaviours of the TF. The optimum number depends on the TF: one with uniform behaviour can be characterised with a smaller set of motifs than one with variable binding affinity, for example.

In more detail, CB-MAR is implemented as follows. Given a TF with a collection of motifs *M* of size *n* and using Tomtom’s Euclidean distance (ED) for motif comparison, we define Pairwise Similarity Score (*PSS*) based on Tomtom P-value *P*_*M*_*i*__. The *PSS* for motif *M*_*i*_ and *M*_*j*_ is computed as:

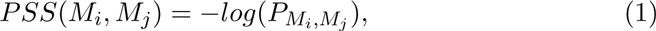
and then normalized by the maximum score of all *PSS* scores of *M*_*i*_. The Average Similarity Score (*ASS*), which we use as the measure of quality and rank, of motif *M*_*i*_, is then computed as:

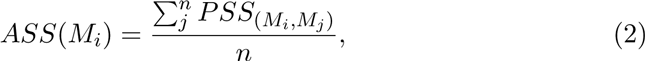

### MARS Web server Implementation

The MARS web server is implemented in Django, a Python web framework, and hosted on an Apache web server while the PWM motifs and sequence benchmark data are stored in a MySQL database. MARS is designed to allow the users to either retrieve ranked motifs for a given TF or rank their own, as long as the required test data is available or uploaded. A guided search function for the available motif and benchmark data assists the users when choosing the tools to use, based on the decision flow diagram in Figure 1. The MARS documentation also provides a detailed guide on the accepted data formats and best practices when using the various tools in the web-server (www.bioinf.ict.ru.ac.za/documentation). For any analysis, the TF name is the only required input, used to retrieve the available motifs from the database and benchmark data in the case of assess-by-score and enrichment methods (Figure 2). Where the data is not available, the user is prompted to upload. MARS currently accepts motifs in MEME format and test ChIP-seq data in BED format.

**Figure 1.**
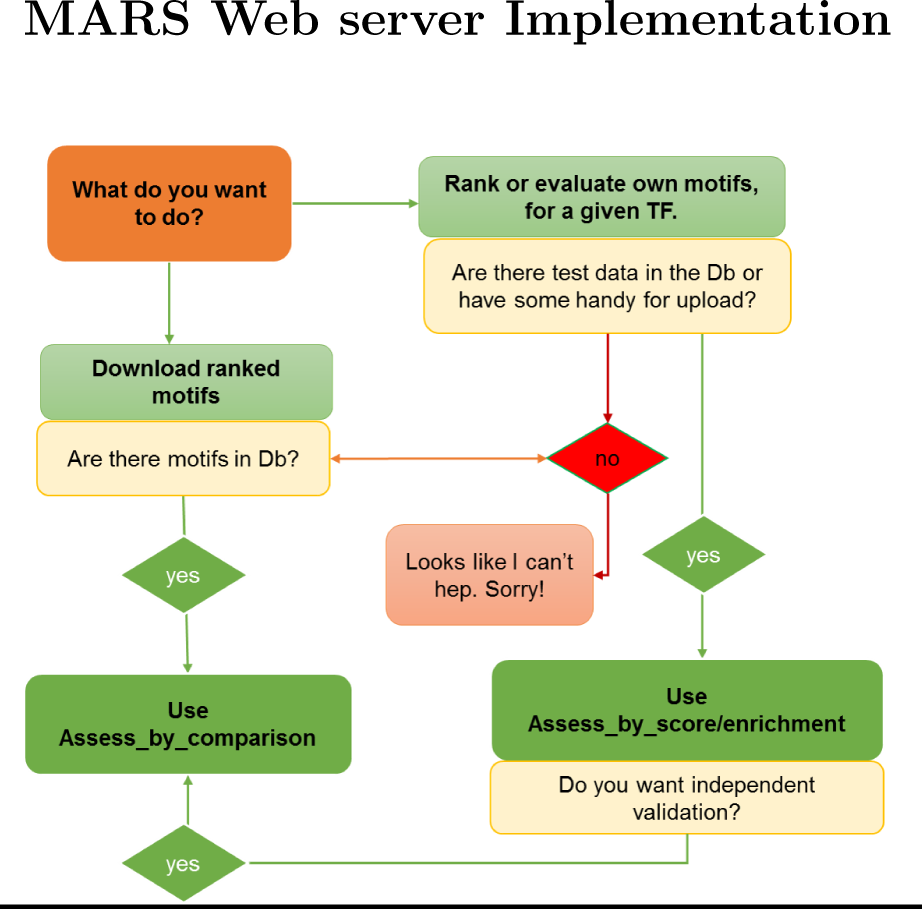
Decision flow diagram. Guides the user on the appropriate tools to use in MARS:

**Figure 2.**
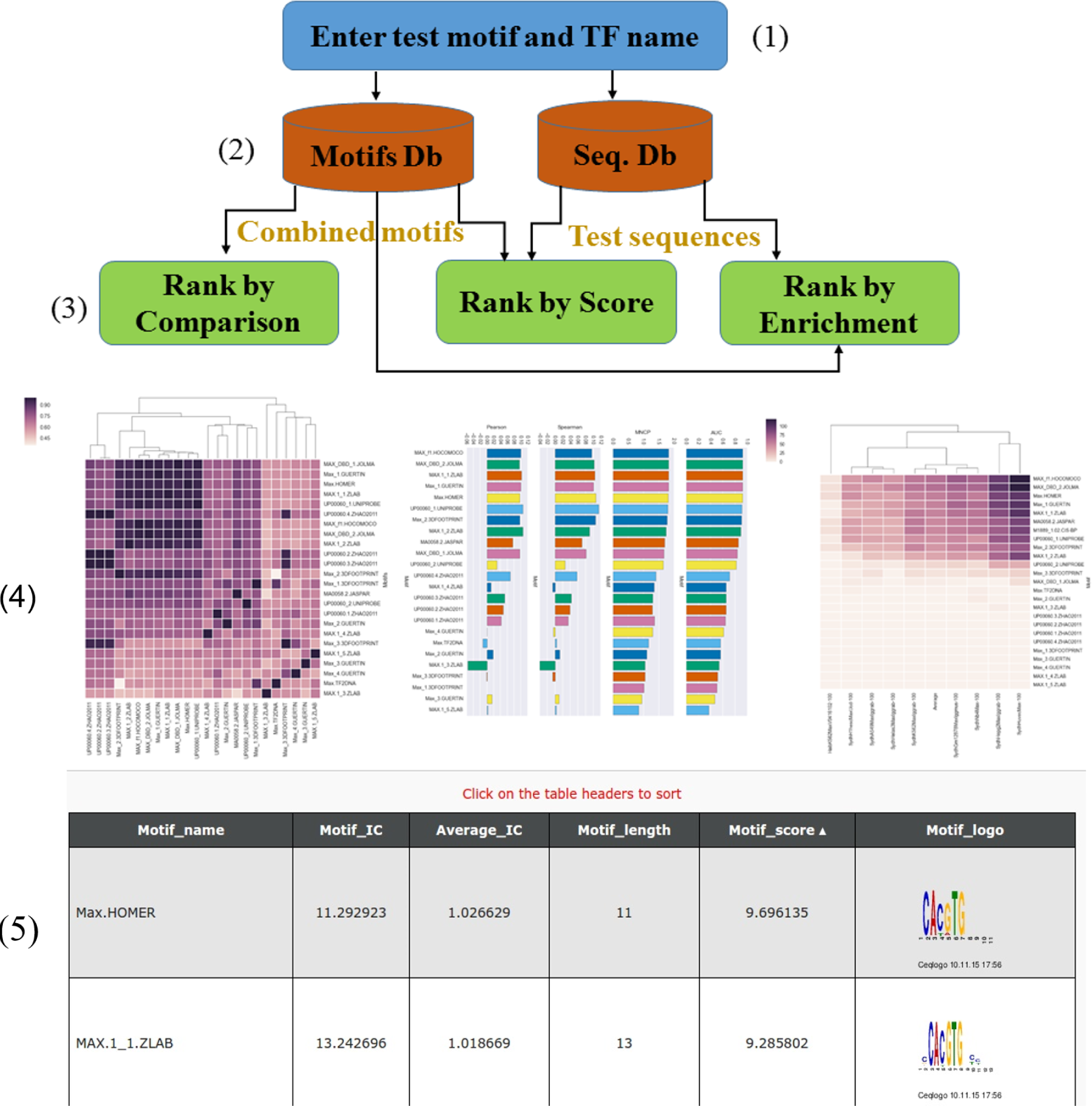
Motif assessment flow diagram: (1) User enters a TF name and/or uploads motifs in MEME format (to evaluate own data). (2) Motifs and test sequences linked to the TF are extracted from the database. (3) Motifs can be ranked, by comparison, used to score test sequences (rank by score) or its enrichment determined using CentriMo (rank by enrichment). (4) The results are visualized interactively, (5) with additional information like motif length, Information content, and logo. The clustergram offers additional details on the motif or test data clustering. In the end, the user can download ranked motifs in MEME format, as well as raw data for further analysis.

## Evaluation of MARS tools

### Comparison of the assessment approaches

How well tools implementing different algorithms and data reproduce each other can act as a crude evaluation. For our evaluation, we select a total of 127 TFs that have a TFClass ID, have more than 10 motifs, and have benchmark data sourced from either PBM (60 TFs) or ChIP-seq (83 TFs) (see Table 1) to rank all the available motifs for each TF using the different tools available. For simplicity of analysis and comparison, we use energy scoring function and AUC statistics throughout these evaluations (Table 2) – a combination we found [18] to produce consistent rankings and is least biased by motif length and information content (IC).

### Motif assessment in *ab initio* motif discovery

To validate CB-MAR, we apply it to choose the best motifs in *ab initio* discovery, a task commonly accomplished using held out data. We take advantage of an ensemble motif finding tool, GimmeMotifs [34], which performs *ab initio* motif discovery from ChIP-seq data using nine algorithms. A total of 110 TFs, which had ENCODE ChIP-seq data and a corresponding TF-class ID, were chosen for the analysis. From all the available data from different cell lines for a given TF, we extract the top 500 peaks widened around the peak centre to 100 bp, merged and shuffled to avoid sampling bias when partitioned. We then perform *ab initio* motif discovery on 50% of the data and the rest is held out for validation. We randomly sampled 5000 peak sequences for Ctcf as it had a large data set to reduce computational costs. After motif discovery, we use our CB-MAR (using Tomtom) approach in combination with motif clustering (using *gimme cluster* from GimmeMotifs at 95% similarity) to rank and narrow down the motif predictions to the best three non-redundant motifs. Next, we use the validation data to evaluate the best motifs identified by CB-MAR and GimmeMotifs using the *gimme roc* command. Internally, GimmeMotifs uses 20% of the input sequences for discovery and the rest for evaluation and ranking.

## Results

### Benchmarking and evaluation of algorithms

For the assess-by-score approach, we have previously performed a thorough comparison and testing to determine the effect of the scoring functions and motif characteristics (length and IC) on score and rank [18]. In summary, we found that motif ranking is influenced by the scoring function used in a TF-specific manner with energy scoring producing the most 'biologically relevant' rankings and that motif IC is not a reliable predictor of motif quality (Table 2). Since there is no ‘ground truth’ by which we can evaluate these motif assessment approaches, we rely on how well rankings from the different tools agree and how well our benchmarks and evaluations meet the requirements listed in the introduction.

For assess-by-comparison, we determine whether CB-MAR is influenced by motif length or information content, which could create a bias, and see if that could explain the difference in performance between Tomtom and FISim. Tomtom scores have a positive correlation (R=0.24, favouring higher IC motifs) with average IC normalized over motif length while FISim scores have a negative correlation (-0.11, penalize higher IC motifs). FISim is not influenced by length (R=0.078) while Tomtom penalizes longer motifs (R=-0.25) since average IC is lower for longer motifs with low IC flanks. Surprisingly, the number of motifs for the TF seems to negatively affect motif scores in FISim (R=-0.38) – due to higher IC as the number of motifs is increased – but has no effect for Tomtom (R=0.011), possibly explaining their difference in performance (Figure 3B).

**Figure 3.**
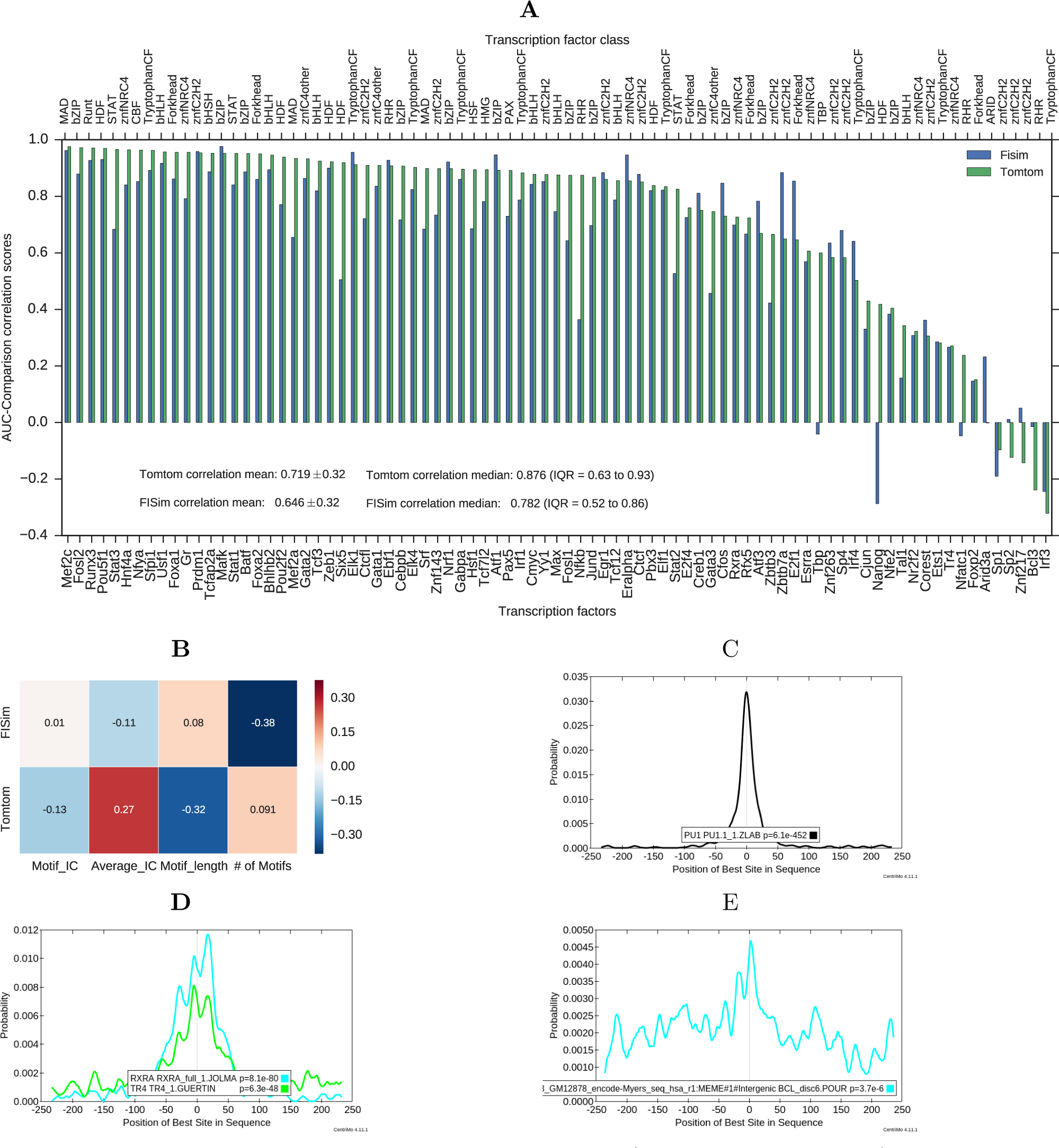
Correlating consistency-based assessment (Tomtom and FISim) with Energy AUC ranking in ChlP-seq data: The bar graph **(A)** shows how rankings based on Energy scoring correlate with consistency based techniques. The mean±STD and median with interquartile range statistics are annotated on the graph. In **(B)**, we show the effect of motif information content (total and averaged by length), the number of motifs (size) and length on motif ranking. The CentriMo plots predict the possible direct binding behaviours (based on the sharp, centred peaks) of **(C)** Pul motifs in ChIP-seq peaks, and indirect or cooperative binding of **(D)** Tr4 and **(E)** Bcl3 motifs. The motif names and the *p*-value of central enrichment of the ChIPed motifs is provided in the figure legends. For Tr4, the other centrally enriched motif (Rxra) could bind cooperatively with it.

### Tomtom comparison produces 'biologically relevant' rankings

For consistency-based motif ranking (CB-MAR), we decide on the best motif comparison algorithms that generate biologically relevant rankings – as defined by how well the motif ranks reproduce those based on *in vivo* data (ChIP-seq)-by correlating with ranks based on energy scoring. From figure 3A, we observe that the scores and ranks based on Tomtom (median=0.88; Interquartile range, IQR=0.63−0.93) can better reproduce AUC scores based on energy scoring compared with FISim (median=0.78; IQR=0.52−0.86). We use the median to summarise the performance of the two techniques since the correlation scores were skewed (Figure 4). The level of correlation between CB-MAR and energy AUC rankings seem to also predict the binding behaviour of the TFs. For Tomtom, we find highly correlated motifs also have centrally enriched peaks in CentriMo (Figure 3B and C) while less or negatively correlated TFs have broad peaks (Figure 3D and E), a known predictor of indirect or cooperative binding [4]. The most common poorly correlated TF family, znfC2H2, are also known to bind in a sequence independent manner [16].

**Figure 4.**
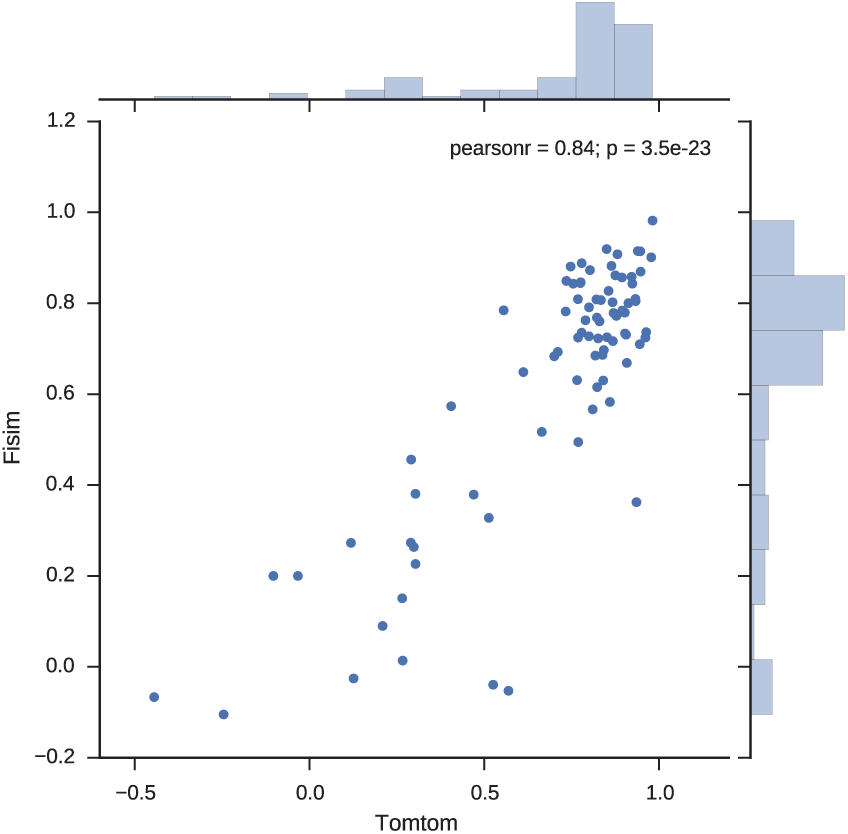
Joint scat-ter plot and histogram. shows the skewed distribution of Spearman rank correlation scores of Tomtom and FISim with those based on Energy scoring

For in vitro data (PBM), however, we do not find a clear performance difference between the median average AUC scores in Tomtom (0.70) and Fisim (0.72), but observe a higher mean in FISim (0.56) compared with Tomtom (0.52), hinting that FISim could better model *in vitro* while Tomtom models *in vivo* binding better. We also note that the TFs with low correlation between scores are known to bind indirectly or cooperatively. Specifically, the TFs from high mobility group (HMG) which have a negative correlation, are known to bind both directly and cooperatively, but they may have a different binding behaviour *in vivo* and *in vitro*. Finally, for TFs with data from both ChIP-seq and PBM (21 TFs), we compared how the Energy scores and motif rankings in the two data types correlate. We observe a similar trend, where the correlation scores reflect the TF binding behaviour (Figure 5). Zbtb3 has a negative correlation of over −0.7, a possible indicator of a difference between *in vivo* and *in vitro* binding behaviour.

**Figure 5.**
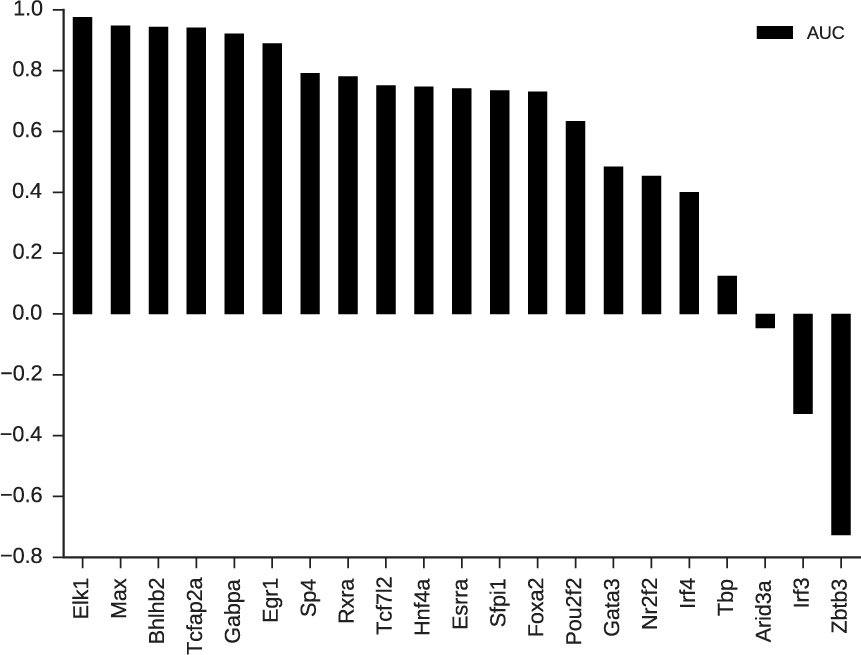
Zbtb3 binds differently *in vivo* and *in vitro*: Motif ranks are in ChIP-seq and PBM benchmark data are mostly in agreement except for a few TFs.

### Assess-by-score-Energy reproduces *gimme roc* rankings

GimeMotifs [34], an ensemble motif discovery pipeline for ChIP-seq data, also includes *gimme roc* for motif quality analysis and ranking. We use this to benchmark our approach and found that *gimme roc* produces motif rankings significantly correlated with the Energy scoring ranks (R=0.99 Pearson, p=1.9 × 10^−105^) (Figure 6) and (**R=0.995 Spearman’s, correlation** p=1.7 × 10^−108^). Also, there is no significant difference between the two sets of scores (**p=0.825, Wilcoxon rank-sum test**), which validates our implementation of energy scoring function.

**Figure 6.**
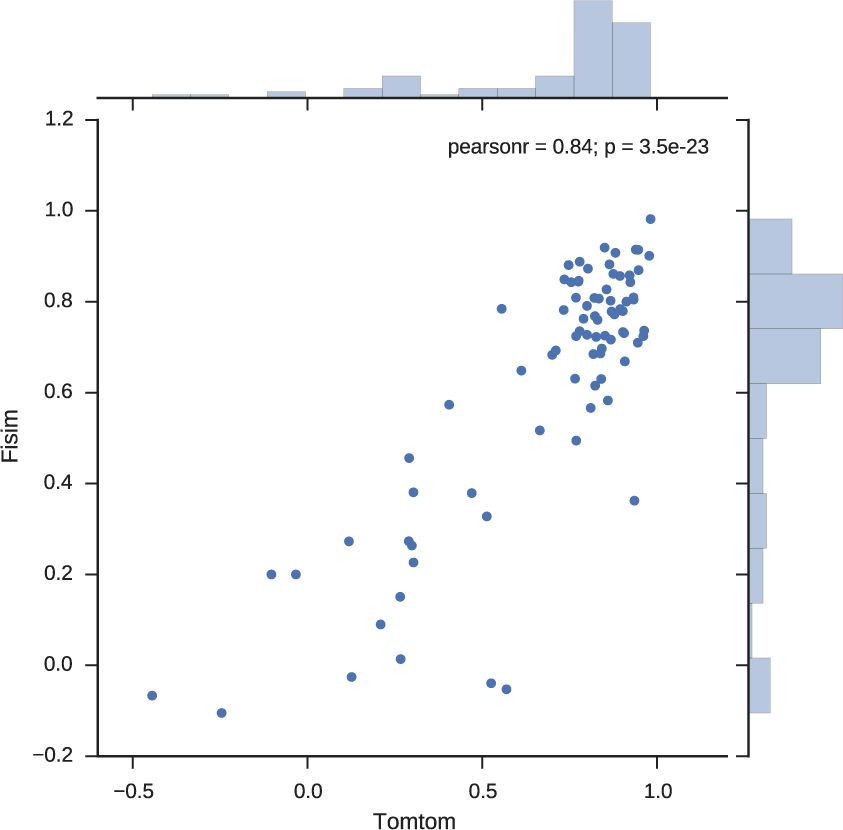
**Joint scat-ter plot and histogram** for *gimme roc* and Energy scoring correlation of AUC scores. The two approaches are in agreement and the data is normally distributed

### CB-MAR generates relevant rankings in motif discovery

The first level of application of motif evaluation is in *ab initio* motif discovery, where an algorithm has to narrow down the identified motifs to a few that reflect the binding behaviour of a TF. In addition, the advent of ensemble motif discovery pipelines makes proper motif assessment and ranking even more desirable. By purely using CB-MAR, we were able to correctly identify better or similar motifs (Figure 7) in a majority of the cases. Overall, the quality of motifs identified by GimmeMotifs and CB-MAR are not significantly different (0.97; Wilcoxon rank-sum test) with mean AUC scores of 0.76 and 0.75 respectively. For 6 TFs that had motifs identified by GimmeMotifs being better than CB-MAR motif by more than 0.1 AUC, we checked if choosing the second or third best motif would have any effect on the quality (Figure 8A). We find that choosing the second motif improves the quality in Maz, Yy1, and Atf1, while the third motif is always of a lower quality except for Irf3. For E2f4, the quality is reduced and no effect is observed on Tcfap2 (Ap2). On the other hand, there were 11 TFs with better scores in CB-MAR than GimmeMotifs. When we choose the second or third motif as ranked by GimmeMotifs the quality of the motifs improved for 7 TFs but no effect in E2f1, Ikzf1, Znf263 and Atf3 TFs (Figure 8B).

**Figure 7.**
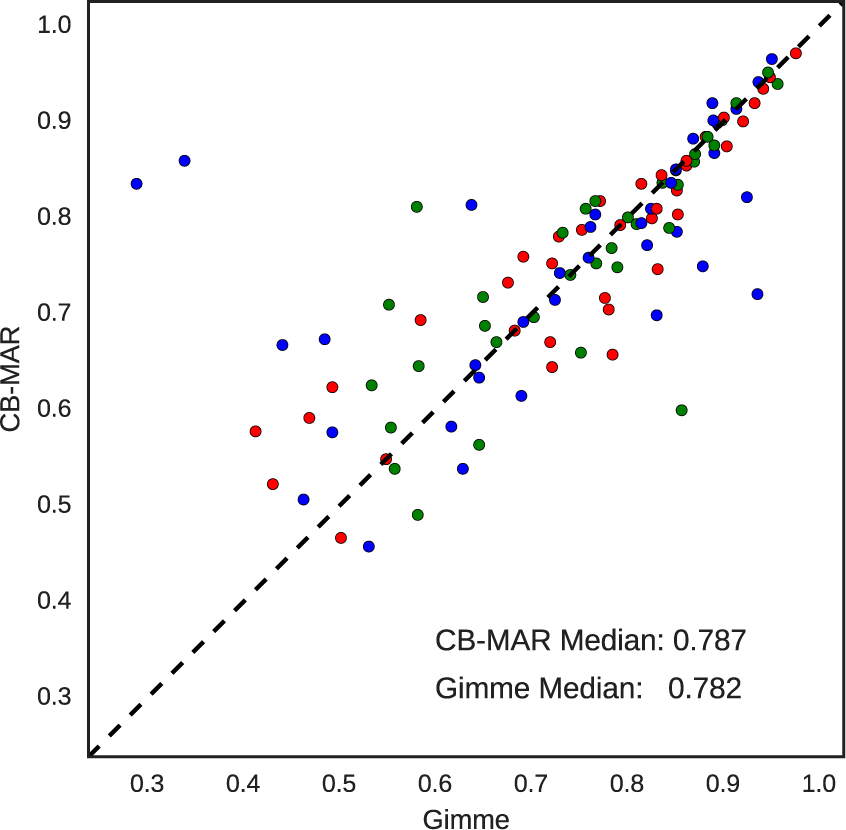
CB-MAR ranking useful in motif discovery: The scatter plot compares the performance of motifs identified by GimmeMotifs and CB-MAR:Tomtom (compare) as evaluated in ChIP-seq data showing the usefulness of data independent approach.

**Figure 8.**
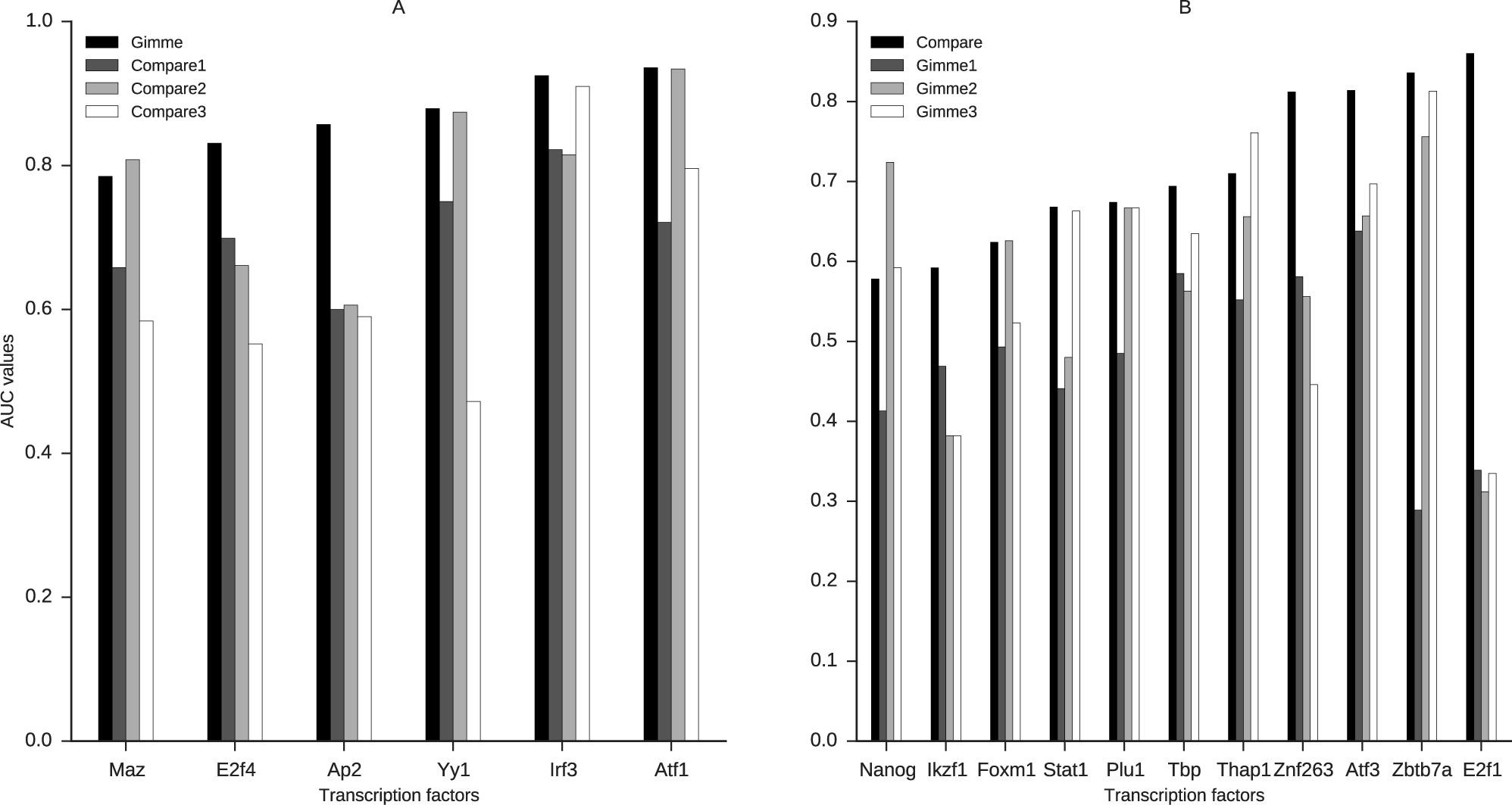
Selecting top three motifs by Gimme and CB-MAR:Tomtom leads to better motifs: The chosen motif is represented by the suffix (Compare1 or Gimme1, for the best and so on). The Y-axis is the mean AUC scores of the motifs in the validation sequences)

## Discussion

The number of motifs available for a single TF continues to increase. This offers variability by increasing the binding spectra captured; TFs bind to degenerate sites spread in the genome. However, this is also a challenge. Choosing a binding model is now a daunting task, given that we can already have up to 47 different PWM models in our collection for a single TF generated from a variety of data and algorithms. How can they be ranked to obtain generalized or specific models for a given task? To address this gap, we introduce MARS, a web server that makes PWM motif evaluation and ranking techniques *accessible*, supported by a database of benchmark data and PWM models. How MARS meets Aniba’s criteria is highlighted in *italics* – see introduction for details. We ensure MARS can *scale* and *evolve* to support new data and algorithms via the modular design of the algorithms and also by allowing users to upload their own benchmark data. Additionally, CB-MAR allows for evaluation *independence*, and we have demonstrated its *relevance* in motif discovery. Given that there is no agreed standard of motif ranking, we offer the end user a variety of techniques and data for their evaluations to ensure *relevance* to different evaluation conditions.

### Motif quality analysis requires systematic comparative assessment

The TF binding spectra are quite diverse. Therefore, a comparative approach to motif evaluation, using a variety of data and techniques, is necessary in order to understand and make an informed decision on motif quality. For example, we can identify difference in binding behaviour of Zbtb3 TF *in vivo* and *in vitro* by correlating motif performance in PBM and ChIP-seq data (Figure 5); Zbtb3 is known to recognize unmethylated motifs *in vivo* and methylated ones *in vitro* [6]. Furthermore, we can discover that HMG TFs may also bind indirectly, cooperatively or a variation of binding behaviour as captured in PBM data by a low or negative correlation between CB-MAR and energy scoring derived ranks (Figure 9). Indeed HMG TFs, specifically the SOX-related factors, are known to bind cooperatively with partner TFs [17,19]. In fact, they are believed to form complexes with partner proteins before recognizing the binding site [17]. Therefore, it is expected that the predicted models in PBM data would differ from how they bind *in vivo*. These observations demonstrate the need for a systematic approach to motif quality assessment.

**Figure 9.**
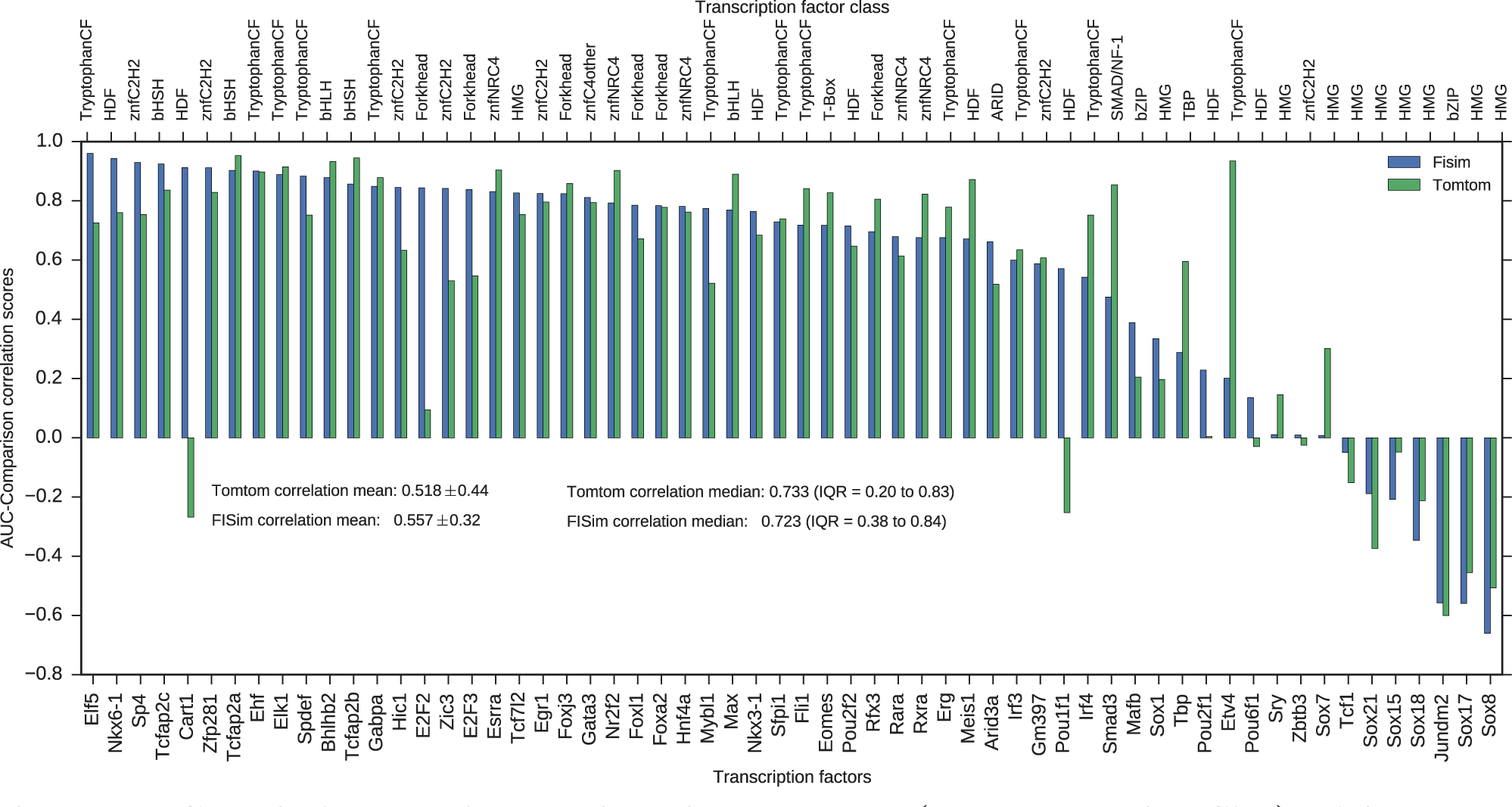
Correlating consistency-based assessment (Tomtom and FISim) with Energy AUC ranking in PBM data: Testing how FISim and Tomtom correlate Energy ranking in PBM data. The TF family class is given on the top axis. The mean ±STD and median with interquartile range statistics are annotated into the figure

### CB-MAR ranks capture *in vivo* binding behaviour

The data-free evaluation approach, CB-MAR, provides a quick and unbiased motif evaluation alternative, especially when the data used in motif discovery is also used for benchmarking. CB-MAR ranks are better correlated with those based on ChIP-seq (R=0.88) than PBM (R=0.73) data, revealing its capability to capture *in vivo* binding (Figures 3A and 9). We further support this argument by using it to successfully identify best PWM models in motif discovery. More details on this in the next section. This implementation reduces the ‘reference motifs’ bias, an approach in which users considered an algorithm successful if it can predict motifs similar to those in a ‘reference database’ at a given (usually arbitrary) similarity threshold. The current collections of ChIP-seq and PBM data in our database can only facilitate data-based quality evaluation for less the 300 TFs out of 1352 that have motifs in our database. This demonstrates that this approach is even more desirable.

Between the two motif comparison algorithms tested for CB-MAR, we show that Tomtom captures in vivo motif ranks–using ChIP-seq data – better than FISim. FISim is designed to favour similarity of high information or conserved sites [11], revealing that scoring high IC sites better may not match biologically similar motifs. Besides, low information flanking sites have been reported to increases binding specificity in some TFs [10,16,21].

### CB-MAR ranking useful in motif discovery

The first step after motif discovery is to filter and narrow down to significant motifs. Usually, a partition of the data is held out for testing, but when there is limited data this may not be feasible. Besides, this is only available to the algorithm developers and to motifs generated using sequencing or microarray data (e.g. promoter sequences, ChIP-seq and PBM) and not to those from TF tertiary structures like 3DFootprint [9]. We have demonstrated that the top performing motifs can be identified by CB-MAR in combination with motif clustering to avoid motif duplicates. However, we do not average similar motifs as done by GimmeMotifs. Rather, in a given cluster, the best ranking motif (by similarity to the rest) is chosen. Motif averaging may produce a motif that does not fit biology or reflect TF binding behaviours as demonstrated by the cases where the GimmeMotifs identified motifs performed significantly worse than CB-MAR (Figure 8B) – evaluated against ChIP-seq data.

## Conclusions

We have developed MARS, a web server hosting a suite of tools for comparative analysis available from www.bioinf.ict.ru.ac.za. This offers choice and flexibility to users since additional test data and motifs can be uploaded, and we do not impose an assessment approach to the users. A major contribution to motif evaluation in this study is the data-independent consistency-based approach (CB-MAR), which offers a good alternative in the absence of benchmark sequence data. We believe that this web server and the algorithms implemented will help reduce motif redundancy and the continued dependence of low quality ‘reference motifs’ due to lack of evaluation data. Our suite also acts as a hub for the motifs collated and annotated from various databases and publications.

## Acknowledgements

The financial assistance of the South African National Research Foundation (NRF) towards this research is hereby acknowledged. Opinions expressed and conclusions arrived at, are those of the authors and are not necessarily to be attributed to the NRF. PM funding: NRF/IFR Grant 85362; CK: DST Innovation Doctoral Scholarship Grant 89071. Finally, we acknowledge the ENCODE Consortium for making the data available.

